# Time-step restrictions for numerical approximations of the Poisson-Nernst-Planck (PNP) equations

**DOI:** 10.64898/2026.04.30.721819

**Authors:** Karoline Horgmo Jæger, Aslak Tveito

## Abstract

The Poisson–Nernst–Planck (PNP) system is an accurate model of electrodiffusion of ionic species. It is commonly used in situations where nanoscale resolution is required, for instance close to ion channels in the membranes of biological cells. The inherent stiffness of the equations has made them challenging to solve and has limited the applicability of the system. In particular, the time step required for stable solutions has typically needed to be very short (nanoseconds), which makes simulations on the time scale of an action potential (milliseconds) difficult. Recently, it has been observed that avoiding operator splitting and instead solving the concentration equations and the electrostatic equation in a coupled manner relaxes the time-step limitation considerably. However, no theoretical explanation of this observation has been provided. Here, we aim to explain why the coupled scheme allows much larger time steps. We illustrate the mechanism by considering special cases that define necessary, but not sufficient, conditions for stability. We also show that these conditions remain relevant for the fully coupled PNP model in 3D.

## 1 Introduction

In biological cells, and in the narrow spaces surrounding them, ion transport is governed by the combined action of diffusion and electrical drift. At larger spatial scales, these effects are often represented by compartment models or by diffusion-based continuum models, and such approximations have been very useful in applications. However, in very small extracellular or intracellular domains, such as the synaptic cleft, the dyad of cardiomyocytes, or the immediate vicinity of ion channels in cell membranes, steep gradients in both concentrations and potential occur over nanometer distances. In such settings, the Poisson–Nernst–Planck (PNP) system provides the natural continuum model of electrodiffusion [1, 2, 3, 4, 5, 6, 7].

Although the PNP system is conceptually straightforward, it is difficult to solve numerically in biologically relevant settings. The equations are strongly coupled and nonlinear, and the formation of thin Debye layers near membranes introduces both steep spatial gradients and very fast temporal dynamics [8, 4]. In practice, this has often implied that very fine spatial grids and extremely small time steps are required in order to obtain stable numerical solutions. As a consequence, direct simulations of the full PNP system on physiological time scales have remained challenging.

In [6], the full PNP equations were solved using an operator-splitting strategy. That approach made it possible to compute informative solutions, but it also led to a severe time-step restriction. The time step used was Δ*t* = 0.02 ns. In [7], this restriction was relaxed considerably by replacing the split treatment by a coupled implicit formulation. The resulting scheme leads, at each time step, to a large but linear coupled system consisting of one elliptic equation for the electrical potential and a set of parabolic equations for the ionic concentrations. The same numerical strategy was used in [9] to solve the full PNP system in the synaptic cleft, again allowing much longer time steps. The time step used there was a million times longer, namely Δ*t* = 0.02 ms. Although these computations showed that the coupled scheme admits time steps that are orders of magnitude larger than those required by the earlier split formulation, a theoretical explanation of this difference is lacking.

A number of numerical approaches for the PNP system have been developed over the last decades. Early work introduced three-dimensional (3D) solvers for ion channel problems and demonstrated that full PNP computations are feasible in realistic geometries [10]. Later, finite-volume and finite-element formulations were developed for biologically relevant electrodiffusion problems and for biomolecular systems with localized fixed charges [1, 11]. Higher-order and structure-preserving schemes have also been proposed [12, 13]. However, despite these advances, the numerical solution of the PNP system remains challenging, and in practice the need for very small time steps may become a severe computational limitation.

The purpose of the present paper is to clarify why the time-step restrictions differ so strongly between these schemes. We first consider a simplified one-dimensional (1D) setting designed to isolate the temporal mechanism responsible for the stiffness. In this setting, we briefly recall the Debye length and the dielectric relaxation time, both of which are standard characteristic scales for electrodiffusion near charged interfaces; see, e.g., [4, 3]. Based on this reduced model, we analyze two numerical schemes:

**Scheme A:** For an operator-splitting scheme of the type used in [6], in which the concentrations are treated implicitly while the electric field is evaluated at the previous time step, we derive a nanosecond-scale time-step restriction ensuring that the discrete solution remains in an invariant region.

**Scheme B:** For the fully coupled implicit scheme of the type used in [7, 9], we show unconditional stability in the sense that the invariant region is preserved for all positive time steps.

We illustrate these theoretical findings by numerical examples in one, two, and three spatial dimensions. In the 3D case, we use the full PNP simulations from [9]. These examples show that the stability properties predicted by the reduced model are also reflected in the corresponding schemes for the full PNP system. In this way, the simplified analysis helps explain the behavior observed in [6, 7, 9], although a theoretical stability analysis of the full 3D PNP system remains out of reach.

## 2 Methods

We consider the Poisson–Nernst–Planck (PNP) model for the electrical potential and the ionic concentrations. This model is given by [14],

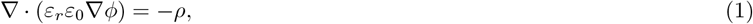

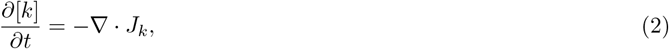

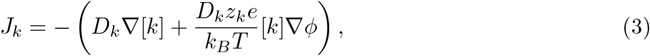

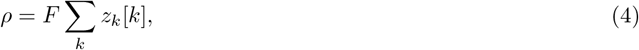

for the considered ion species, *k*, where [*k*] is the concentration of ion species *k* (in mM), *φ* is the electric potential (in mV), *J*_*k*_ is the flux of ion species *k* (in mMnm/ms), and *ρ* is the charge density (in C/m^3^). The quantities *ε*_*r*_, *ε*_0_, *D*_*k*_, *z*_*k*_, *e, k*_*B*_, *T*, and *F* are model parameters and physical constants listed in Table 1.

**Table 1:** Parameter for the PNP system. Here, Ω_*e*_ refers to the extracellular domain, Ω_*m*_ refers to the membrane domain (including the membrane and mouth of the synaptic vesicle), and Ω_*i*_ refers to the intracellular domain (including the synaptic vesicle).

| Parameter | Description | Value | Ref. |
| --- | --- | --- | --- |
| $F$ | Faraday's constant | 96485.3365 C/mol | [15] |
| $e$ | Elementary charge | $1.60217662 \cdot 10^{-19}$ C | [15] |
| $k_B$ | Boltzmann constant | $1.380649 \cdot 10^{-20}$ mJ/K | [15] |
| $T$ | Temperature | 310 K | |
| $\varepsilon_0$ | Vacuum permittivity | 8854 fF/m | [15] |
| $\varepsilon_1$ | Relative permittivity, $\varepsilon_r$, in $\Omega_e$ and $\Omega_i$ | 80 | [10] |
| $\varepsilon_m$ | Relative permittivity, $\varepsilon_r$, in $\Omega_m$ | 2 | [10] |
| $D_{\text{Na}^+}$ | Default diffusion coefficient for $\text{Na}^+$ | $1.33 \cdot 10^6$ nm<sup>2</sup>/ms | [16] |
| $D_{\text{K}^+}$ | Default diffusion coefficient for $\text{K}^+$ | $1.96 \cdot 10^6$ nm<sup>2</sup>/ms | [16] |
| $D_{\text{Ca}^{2+}}$ | Default diffusion coefficient for $\text{Ca}^{2+}$ | $0.71 \cdot 10^6$ nm<sup>2</sup>/ms | [16] |
| $D_{\text{Cl}^-}$ | Default diffusion coefficient for $\text{Cl}^-$ | $2.03 \cdot 10^6$ nm<sup>2</sup>/ms | [16] |
| $D_{\text{Glu}^-}$ | Default diffusion coefficient for $\text{Glu}^-$ | $0.86 \cdot 10^6$ nm<sup>2</sup>/ms | [17] |
| $z_{\text{Na}^+}$ | Valence of $\text{Na}^+$ | 1 | |
| $z_{\text{K}^+}$ | Valence of $\text{K}^+$ | 1 | |
| $z_{\text{Ca}^{2+}}$ | Valence of $\text{Ca}^{2+}$ | 2 | |
| $z_{\text{Cl}^-}$ | Valence of $\text{Cl}^-$ | -1 | |
| $z_{\text{Glu}^-}$ | Valence of $\text{Glu}^-$ | -1 | |

### 2.1 Numerical examples

In order to compare the stability of the two numerical schemes A and B described above, we consider three different test cases with increasing complexity.

#### 2.1.1 Case I: A simple 1D test problem

As a first test case, we consider a 1D domain of length 10 nm. For the electric potential, we impose Dirichlet boundary conditions,

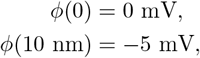

while homogeneous Neumann boundary conditions are imposed for all concentrations. We include the three ionic species Na^+^, Cl^−^, and Glu^−^, with initial concentrations given by the spatially constant values 100 mM, 99 mM, and 1 mM, respectively. The diffusion coefficients are taken from Table 1, and the relative permittivity is set equal to *ε*_1_ throughout the domain. The spatial discretization is uniform with mesh size Δ*x* = 0.2 nm.

#### 2.1.2 Case II: A simple 2D problem with a membrane

In the second test case, we consider two parts of a two dimensional (2D) domain separated by a membrane. On each side of the membrane, we include the two ionic species, Na^+^ and Cl^−^. We set the initial condition of Na^+^ to 100 mM on both sides of the membrane and the initial condition of Cl^−^ to 100.1 mM and 99.9 mM on the left and right parts of the membrane, respectively. We use homogeneous Neumann boundary conditions for both *φ* and the concentrations on all boundaries, except for the Dirichlet boundary condition *φ* = 0 mV on the left boundary. In the membrane, the diffusion coefficients are set to zero, whereas in the remaining domain they are set to the values from Table 1. We discretize the domain using Δ*x* = 0.1 nm and Δ*y* = 1 nm.

#### 2.1.3 Case III: A 3D synaptic cleft

As a final example we consider a 3D domain representing a synaptic cleft. More specifically, we represent a part of a presynaptic cell with a synaptic vesicle docked in the membrane, a part of a postsynaptic cell, the membranes of the cells, the extracellular cleft between the cells, and the remaining extracellular space surrounding them. We use the same setup as in [9], but because we need to take small time steps to ensure stability of scheme A, we only consider the dynamics in the first hundred nanoseconds following release from the synaptic vesicle. This is too early for the AMPA receptors on the postsynaptic membrane to be activated. Therefore, we do not include AMPA receptors (or any other postsynaptic receptors) in these simulations.

We consider the five ionic species Na^+^, K^+^, Ca^2+^, Cl^−^ and Glu^−^. In the membrane, all ionic concentrations and diffusion coefficients are set to zero. The initial conditions for the ionic species in the remaining parts of the domain are as given in [9]. In the synaptic cleft between the cells, the default diffusion coefficients are divided by a factor 2.56 to represent reduced diffusion in the narrow space (consistent with, e.g, [17]). The geometry of the synapse and the initial conditions for the ionic species are the same as those applied in [9].

## 3 Results

We begin the analysis of the numerical schemes by considering simplified problems designed to provide insight into the spatial and temporal dynamics of both the analytical solution of the PNP equations and their numerical approximations.

### 3.1 A simplified 1D model

We consider a 1D domain *x* ∈ [0, *L*] with two ionic species: a positive ion *p* (modeled after Na^+^) with valence *z*_*p*_ = +1, and a negative ion *n* (modeled after Cl^−^) with valence *z*_*n*_ = −1. The concentrations *p* = *p*(*x, t*) and *n* = *n*(*x, t*) are given in mM, and the electric potential *φ* = *φ*(*x, t*) in mV. We define

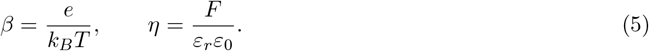

The fluxes of the two ionic species are given by

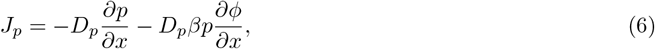

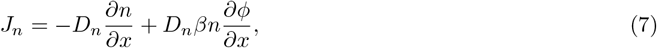

and the 1D PNP system reads

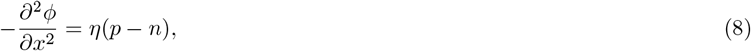

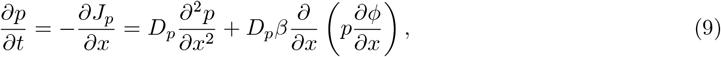

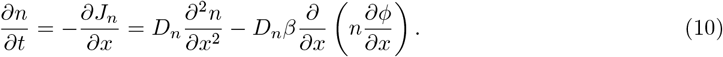

### 3.2 The Debye length in the stationary case

To identify the characteristic spatial scale, we consider the stationary problem on the half-line *x* ∈ [0,∞). In the stationary case the ionic fluxes vanish, and therefore

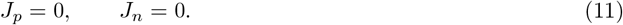

Using the definitions of the fluxes in (6)–(7), this gives

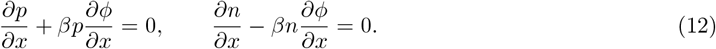

Integrating these equations, we obtain

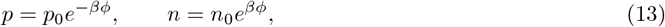

where *p*_0_ and *n*_0_ are constants. Substituting into Poisson’s equation (8) yields the Poisson–Boltzmann equation

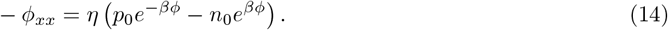

We now linearize around an electroneutral reference state with

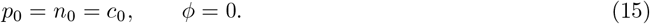

Using the first-order approximations

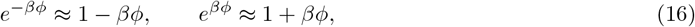

equation (14) reduces to

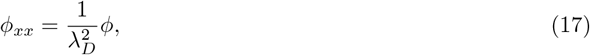

Where

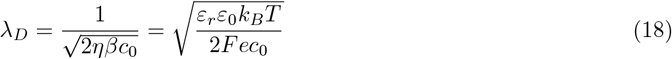

The physically relevant solution is the decaying solution

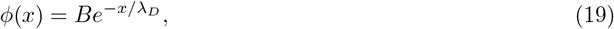

which shows that *λ*_*D*_ is the *e*-folding distance over which the boundary perturbation decays into the bulk. The Debye length therefore defines the natural spatial scale of charge separation in the problem. Using *c*_0_ = 100 mM, *ε*_*r*_ = *ε*_1_ and the parameter values from Table 1, we obtain

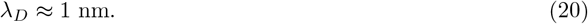

### 3.3 The dielectric relaxation time

To identify the characteristic temporal scale of charge relaxation, we linearize the 1D PNP system around the electroneutral reference state *p* = *n* = *c*_0_, and *φ* = 0. For simplicity, assume equal diffusion coefficients, *D*_*p*_ = *D*_*n*_ = *D*. Writing

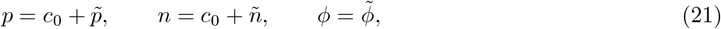

and defining the charge imbalance by

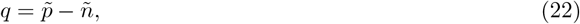

subtracting (10) from (9) gives

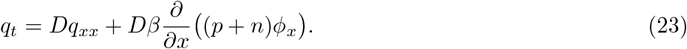

Expanding the last term and retaining only first-order terms yields

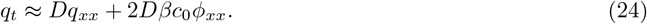

Using Poisson’s equation (8),

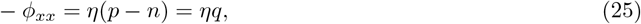

we obtain

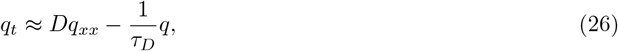

where

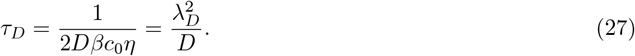

To interpret (26), consider Fourier modes of the form

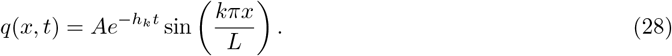

Substitution into (26) gives

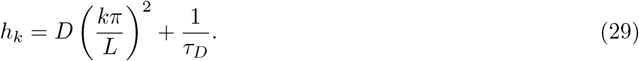

Thus the decay of *q* is governed by both diffusion and dielectric relaxation. Using the parameter values from Table 1, with *L* = 10 nm, *c*_0_ = 100 mM, *ε*_*r*_ = *ε*_1_, and *D* taken as the average of 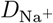 and 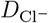, we find

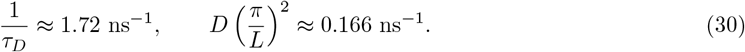

Hence, even for the fundamental mode, the decay is dominated by the dielectric relaxation term, and the relaxation toward electroneutrality takes place on the nanosecond scale.

### 3.4 Temporal and spatial characteristics

The characteristic temporal and spatial scales identified above are on the order of nanoseconds and nanometers, respectively. This means that a numerical scheme aiming to fully resolve the PNP equations in both space and time must, in principle, use spatial and temporal discretizations consistent with these scales. In space, this is not unreasonable, since the applications of interest involve very small domains, often with dimensions of only a few nanometers. In time, however, the situation is different. The physiological processes of interest typically evolve on the microsecond to millisecond scale, and resolving the dynamics with nanosecond time steps is therefore computationally very demanding and may not be necessary. We therefore seek numerical schemes that allow time steps far larger than the nanosecond scale. This was achieved in [9], and our aim here is to clarify why that particular scheme admits such large time steps while remaining stable.

### 3.5 A simplified temporal model

To study the time-step restrictions, we return to the simplified 1D model (8)–(10) and consider a simplified setting in which *p* and *n* are spatially constant. Then *p*_*x*_ = *n*_*x*_ = 0, and the system (8)–(10) reduces to

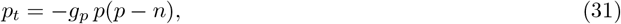

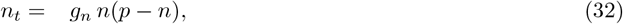

where

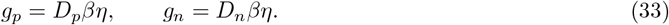

We next introduce two finite-difference schemes for (31)–(32). These schemes are chosen to reflect, in a simplified setting, the temporal treatment used in [6] (Scheme A below) and in [7, 9] (Scheme B below). We derive stability conditions for these schemes for the reduced system (31)–(32) and then demonstrate below that the same conditions are reflected in computations for the full 3D PNP model.

#### 3.5.1 Scheme A: operator splitting

In [6], the PNP system was solved by an operator-splitting strategy: the electrostatic equation (1) was solved first, and the resulting electric field was then used in the concentration equations (2), where the concentrations were treated implicitly. In terms of the simplified temporal model (31)–(32), this leads to the scheme

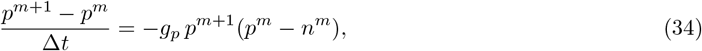

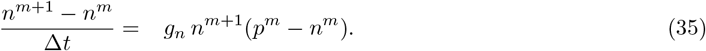

Solving for *p*^*m*+1^ and *n*^*m*+1^ gives

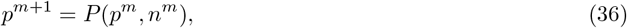

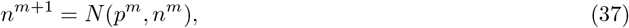

where

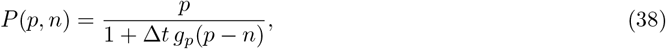

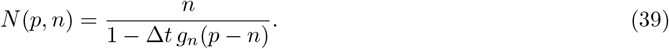

We next show that this scheme preserves the invariant region [0, *Q*] *×* [0, *Q*], where *Q* = max(*p*^0^, *n*^0^). The derivatives of *P* and *N* are

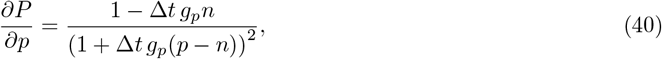

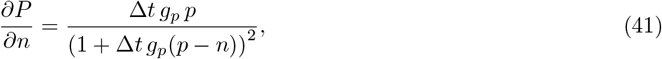

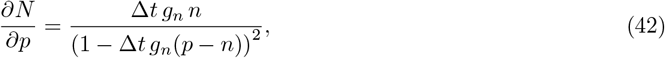

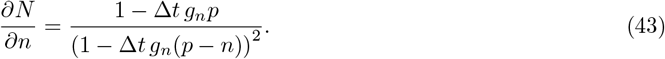

These quantities are nonnegative on [0, *Q*] *×* [0, *Q*] provided

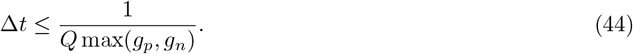

Hence both *P* and *N* are increasing in both variables on the square [0, *Q*] *×* [0, *Q*]. By monotonicity,

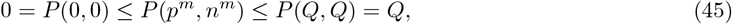

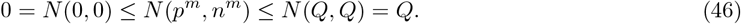

Therefore, by induction,

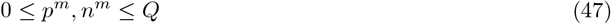

for all *m*, provided (44) holds.

For *Q* = 100 mM, *ε*_*r*_ = *ε*_1_, and the parameter values from Table 1, we get *g*_*p*_ ≈ 6782 (mMms)^−1^ and *g*_*n*_ ≈ 10351 (mMms)^−1^, and condition (44) becomes

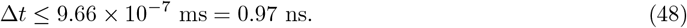

Moreover, using *ε*_*r*_ = *ε*_*m*_ in the definition of *η*, the condition reads

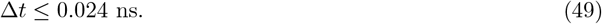

Thus Scheme A requires a time step on the nanosecond scale. This is consistent with the results obtained in [6], and with the numerical experiments presented below.

#### 3.5.2 Scheme B: implicit coupled scheme

It was observed in [7, 9] that by solving the electrostatic equation (1) fully coupled with the concentration equations (2), one could use much longer time steps without observing instabilities. In order to avoid solving a nonlinear system at each time step, the concentration factor in the drift term, that is, the factor multiplying the electric field in (3), was treated explicitly. For the simplified temporal model (31)–(32), this leads to the scheme

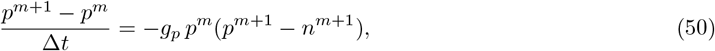

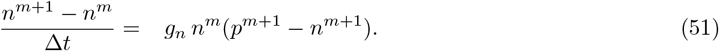

referred to as Scheme B. This scheme may be written in matrix form as

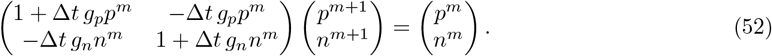

Solving this system gives

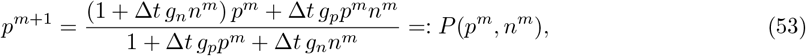

and

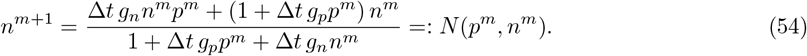

Furthermore,

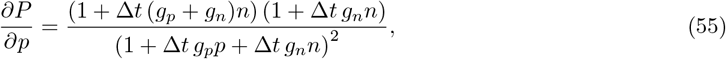

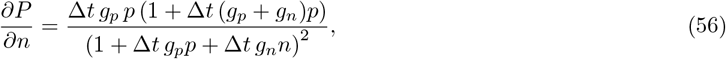

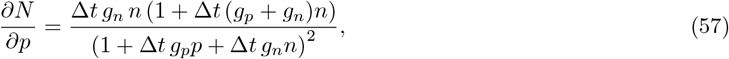

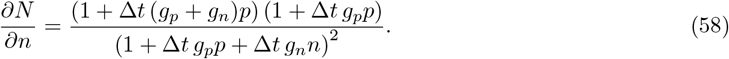

Since all four derivatives are positive for *p, n* ≥ 0, both *P* and *N* are increasing in both variables on the square [0, *Q*] *×* [0, *Q*]. Moreover,

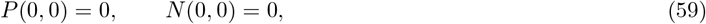

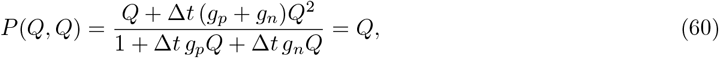

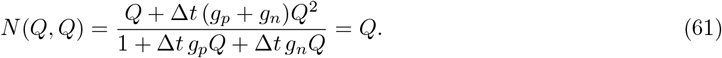

Therefore,

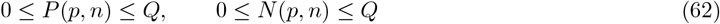

for all (*p, n*) ∈ [0, *Q*] *×* [0, *Q*]. Hence, if

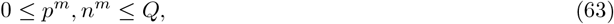

then

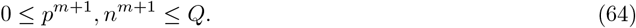

By induction, it follows that

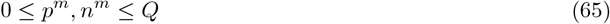

for all *m*.

We conclude that Scheme B preserves the invariant region

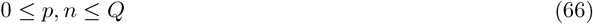

for every Δ*t >* 0. In this sense, Scheme B is unconditionally stable.

### 3.6 Numerical simulations

We now turn to numerical simulations of the full PNP model in order to assess whether the stability bounds derived for the simplified temporal model (31)–(32) are informative also in the full setting.

#### 3.6.1 Case I: Simple 1D test problem

The first numerical example is a simple 1D problem the one studied in Figure 7 of [9]. The set up is described in the Methods section. In short, we consider a domain of length *L* = 10 nm with homogeneous no-flow Neumann boundary conditions for the concentrations. Furthermore, on the left side of the domain, we apply the Dirichlet boundary condition *φ* = 0 mV, and on the right side, we apply the Dirichlet boundary condition *φ* = −5 mV. We include the three ionic species Na^+^, Cl^−^, and Glu^−^, with initial concentrations given by the spatially constant values of 100 mM, 99 mM, and 1 mM, respectively.

In Figure 1, we show numerical solutions of the problem. We consider each of the two numerical schemes as well as different values of Δ*t* and focus on time points when stationary solutions of the problem should have been achieved. We display the solutions in the last time step as a solid black line and the solutions in the second to last time step as a gray dotted line. The second to last time step is included to reveal potential oscillations in the numerical solutions.

**Figure 1:**
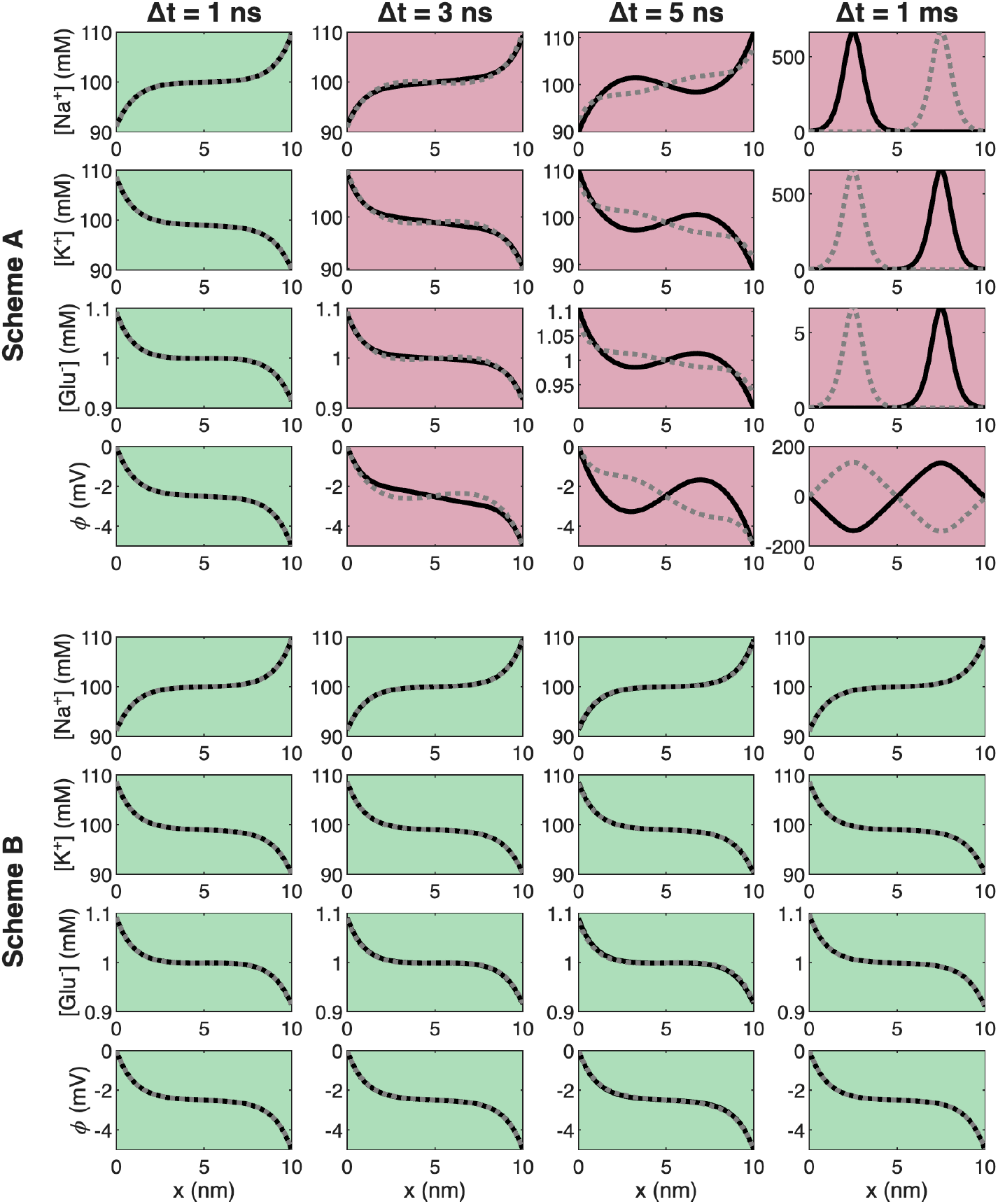
Solutions of a simple 1D test problem for the two considered schemes for different values of Δ*t*. The solid black line is the solution at 20 ns for Δ*t* = 1 ns and Δ*t* = 5 ns, the solution at 21 ns for Δ*t* = 3 ns, and the solution at 20 ms for Δ*t* = 1 ms. In all cases, the dotted gray line is the solution at the second to last time step, included to reveal potential oscillations.

We observe that for Scheme B, the solutions are equal for all the considered values of Δ*t*, and the solutions in the second to last time steps are very similar to the solutions in the last time step, indicating that stationary solutions have been achieved and that the numerical scheme is stable. For Scheme A, on the other hand, the numerical solutions appear to be free of oscillations only for the finest time step considered (Δ*t* = 1 ns). For Δ*t* = 3 ns, small oscillations are visible, and the size of the oscillations increases as the size of the time step is increased.

#### 3.6.2 Case II: Simple 2D test problem with a membrane

Next, we consider a simple 2D test problem with two parts of a domain separated by a membrane, like in Figure 5 in [7]. We include the two ionic species, Na^+^ and Cl^−^, and the initial conditions are set up such that the domain is not electroneutral. More specifically, Na^+^ is initially 100 mM on both sides of the membrane and Cl^−^ is initially 100.1 mM and 99.9 mM on the left (Ω_*L*_) and right (Ω_*R*_) parts of the membrane, respectively. Homogeneous no-flow Neumann boundary conditions are applied for the concentrations on all boundaries and between the membrane and Ω_*L*_ and Ω_*R*_. Therefore, ions cannot leave or enter the two domain parts and electroneutrality cannot be reached during the simulation. Instead, a Debye layer is formed close to the membrane.

In Figure 2, we display the numerical solutions of the problem close to the membrane. The solutions are recorded at a time step when stationary solutions should have been achieved. Again, we plot the last time step using a solid black line and the second to last time step using a dotted gray line to spot potential oscillations. We observe that for a time step of Δ*t* = 0.9 ns, Scheme A provides stable and accurate solutions. However, for Δ*t* = 1 ns, small oscillations are present, and for Δ*t* = 1.5 ns, the oscillations are larger. Moreover, for Δ*t* = 1 ms, the solutions of Scheme A are completely wrong. For Scheme B, on the other hand, the solutions appear to be stable and accurate for all the considered values of Δ*t*.

**Figure 2:**
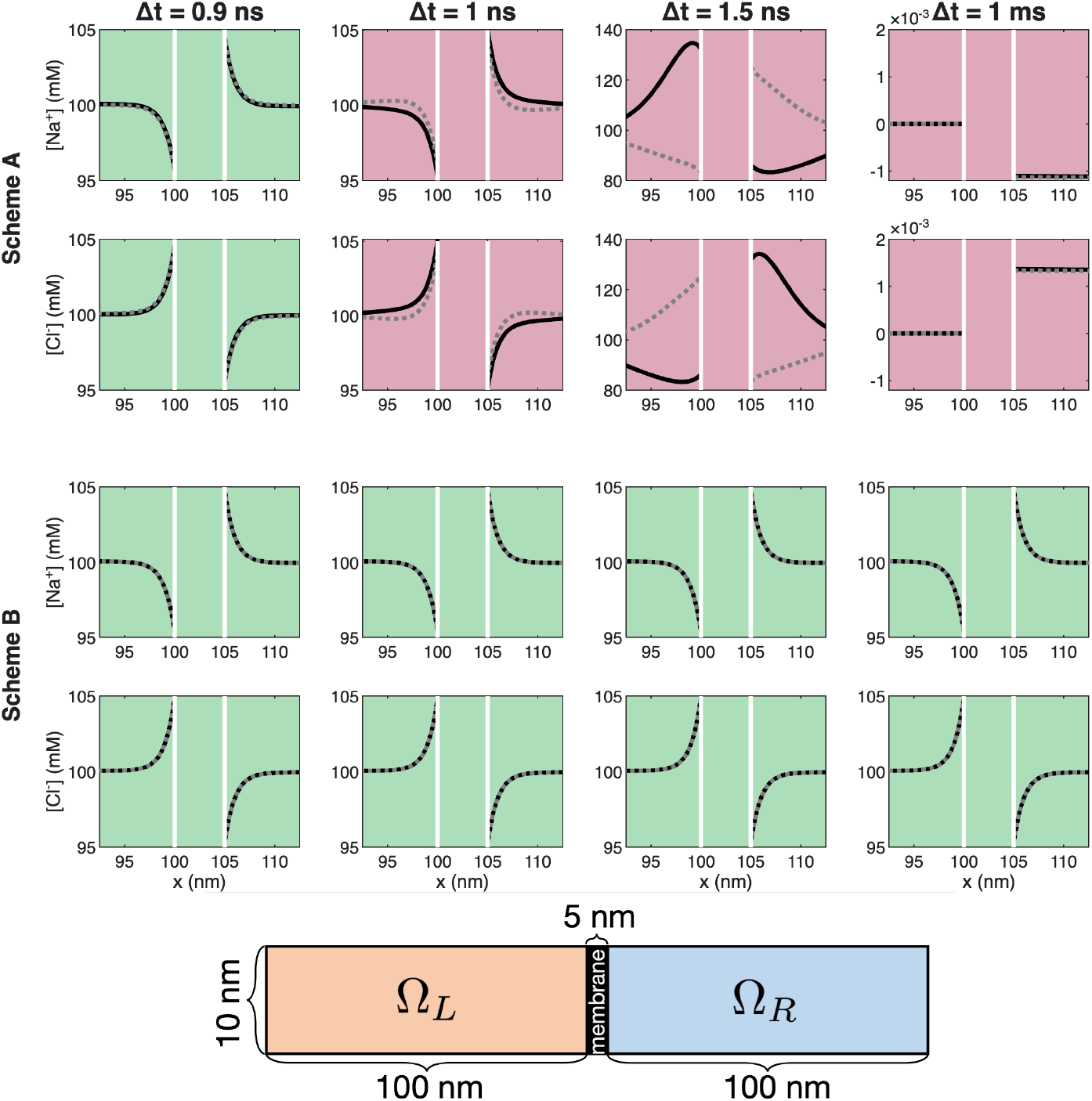
Solutions of a simple 2D test problem with a membrane for the two considered schemes for different values of Δ*t*. We plot the solutions along a line in the *x*-direction. For Δ*t* ≤1.5 ns, the solid black line is the solution at 9 ns, and for Δ*t* = 1 ms, the solid black line is the solution at 9 ms. In all cases, the dotted gray line is the solution at the second to last time step, included to reveal potential oscillations in the numerical solutions. The white vertical lines represent the membrane boundaries. The domain geometry is displayed in the lower panel.

#### 3.6.3 Case III: A full 3D synaptic cleft simulation

Finally, we consider a full 3D synaptic cleft simulation like the one studied in [9]. We focus on the dynamics in the extracellular cleft between a presynaptic and a postsynaptic neuron following the release of the neurotransmitter Glu^−^ from a synaptic vesicle docked at the presynaptic membrane. Figure 3 shows the Glu^−^ concentration and *φ* in a point in the center of the domain as functions of time for the two numerical schemes and different values of Δ*t*. The lower right panels show illustrations of the considered domain.

**Figure 3:**
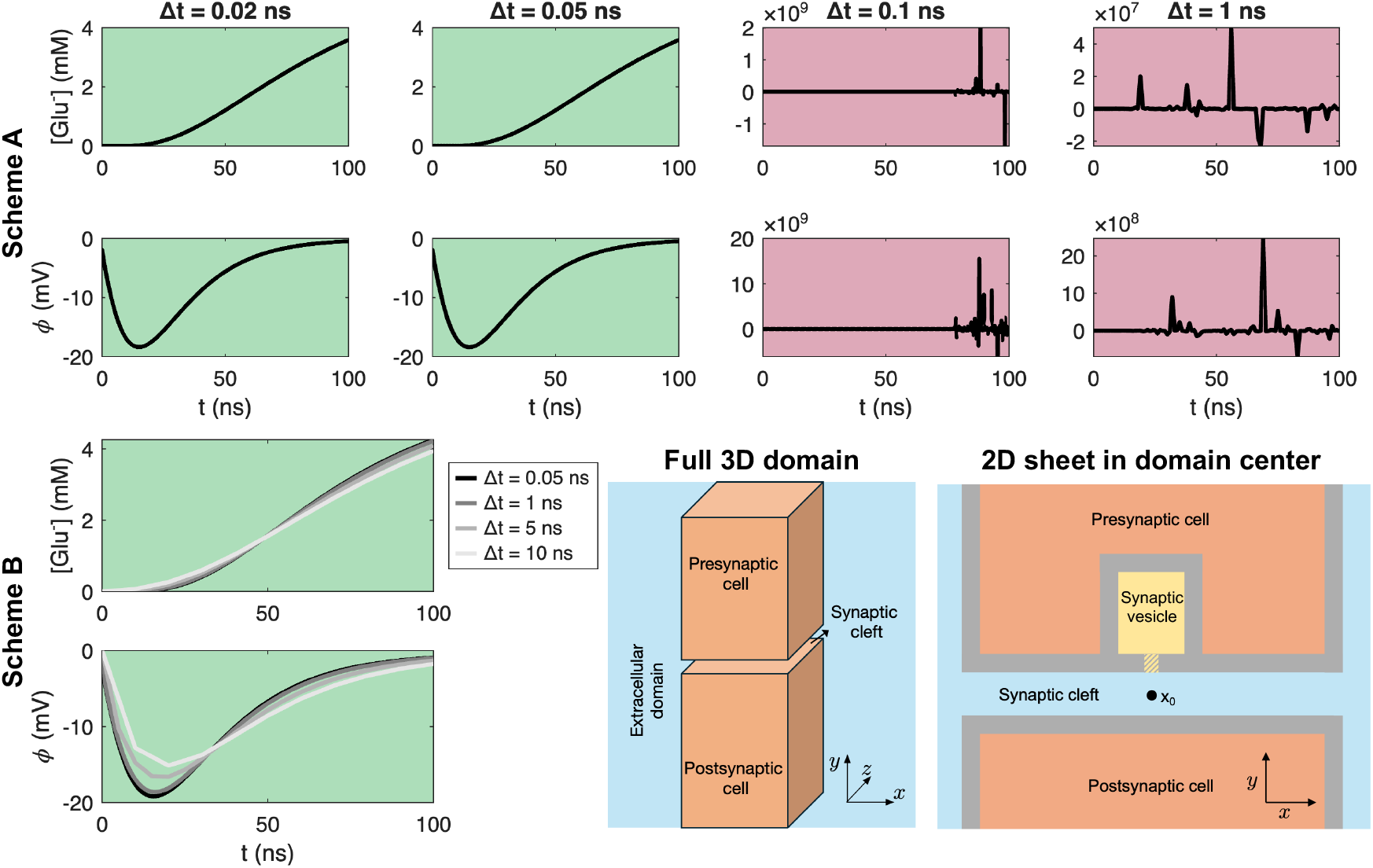
Solutions for the two considered schemes for different values of Δ*t* of full 3D synaptic cleft simulations. The domain setup is illustrated in the lower right panels. When the simulation starts, the mouth of the synaptic vesicle opens allowing Glu^−^ ions to flow from the synaptic vesicle into the synaptic cleft. In the remaining panels, the Glu^−^ concentration and *φ* in the center point marked by *x*_0_ are plotted as functions of time. For Scheme B, the solutions for the different values of Δ*t* are plotted in the same panel to investigate the accuracy of the solutions.

In the stable solutions, we observe that the Glu^−^ concentration in the cleft increases as Glu^−^ is allowed to flow out of the synaptic vesicle. In addition, we observe that a negative electrical potential is generated in the cleft. For Δ*t* = 0.02 ns and Δ*t* = 0.05 ns, Scheme A seems to provide stable solutions, but for Δ*t* = 0.1 ns or Δ*t* = 1 ns, we get wild oscillations in the solutions. Furthermore, as before, Scheme B provides stable solutions for all the considered values of Δ*t*.

In [9], we observed that Scheme B gave stable solutions in simulations of the synaptic cleft dynamics using a time step of Δ*t* = 0.02 ms. For stability, even larger time steps could have been used, but Δ*t* = 0.02 ms was chosen for accuracy at the millisecond time scale. Likewise, in the lower left panels of Figure 3, we observe that even though Scheme B is stable for large values of Δ*t*, a small value of Δ*t* might be required in order to get sufficiently accurate solutions on the nanosecond scale following synaptic vesicle release.

## 4 Discussion

The main observation of this paper is that the severe nanosecond-scale time-step restriction observed for operator-splitting methods is not an unavoidable consequence of the PNP system itself, but rather of how the electrostatic and concentration equations are treated numerically. By replacing the split treatment with a coupled implicit formulation, the restriction is removed in the simplified temporal model, and the same qualitative behavior is observed in the numerical experiments for the full PNP system.

### 4.1 Earlier numerical work

Early work on the PNP equations pointed to the stiffness of the time-dependent Nernst–Planck equations [18]. Later, numerical solvers for the full PNP system were developed in biologically relevant geometries [1, 11], and alternative formulations based on electroneutrality were introduced in order to avoid resolving the Debye layer explicitly [19, 20]. More recently, structure-preserving discretizations with unconditional stability properties have been constructed [13]. The present paper is narrower in scope. Rather than proving stability for the full PNP system, we aim to explain, by a simplified analysis, why the coupled treatment permits much larger time steps than the split formulation.

### 4.2 Origin of the stiffness

The PNP system is a precise continuum model of electrodiffusion and has been used in several biologically relevant settings, including ion transport near cell membranes, in the synaptic cleft, and in narrow extracellular spaces [4, 2, 3, 1, 21, 20]. The source of the stiffness is the tight coupling between the concentration equations and Poisson’s equation. This coupling introduces a spatial scale given by the Debye length and a temporal scale given by the dielectric relaxation time. For the parameter values considered here, these scales are on the order of 1 nm and 1 ns, respectively. The spatial scale is compatible with the applications of interest, since domains such as the cardiac dyad [7], the neuronal synaptic cleft [9], and the intercalated disc of cardiomyocytes [6] all have dimensions on the order of a few to a few tens of nanometers. The temporal scale is more problematic. Physiological processes typically evolve on the micro-to millisecond scale, and a numerical method requiring nanosecond time steps is therefore impractical for many relevant simulations.

### 4.3 Reduced model

To isolate the mechanism behind the time-step restriction, we considered a purely time-dependent reduction of the PNP system. This reduced model is simple enough to permit a complete analysis, while still retaining the essential coupling responsible for the stiffness. For the split scheme, the analysis reproduces the nanosecond-scale restriction observed in computations with the full PNP system [6]. For the coupled scheme, the same analysis shows that the invariant region is preserved for all positive time steps. This explains why the coupled formulation behaves so differently in practice. In particular, it is consistent with the fact that the time step used in [9] is many orders of magnitude larger than the one used in [6], while the same numerical strategy was introduced for dyadic electrodiffusion in [7].

### 4.4 Limitations and practical implications

The analysis has clear limitations. The unconditional stability result is derived only for the simplified temporal model, not for the full PNP system in space. The reduced model provides necessary, but not sufficient, stability information, and a full theoretical analysis of the coupled scheme for the complete 3D PNP equations remains open. The numerical experiments in Cases I–III support the relevance of the reduced model, but they certainly do not constitute a proof.

Stability does not imply that arbitrarily large time steps are acceptable. In the 3D synaptic cleft simulations, the coupled scheme remains stable for large Δ*t*, but accurate resolution of the rapid dynamics immediately following vesicle release still requires small time steps [9]. For slower dynamics on the micro- or millisecond scale, much larger steps can be both stable and accurate [7, 9]. The appropriate choice of Δ*t* must therefore be determined by the time scale of the phenomenon under study, not by stability considerations alone. This suggests that adaptive time-stepping may be useful, allowing small steps during rapid transients and larger steps during slower phases.

### 4.5 Analytical approaches

Analytical approaches to electrodiffusion based on the PNP equations have been pursued for a long time; see, e.g., [22]. The numerical schemes considered here are motivated by the difficulty of solving the PNP equations directly. Analytical progress is limited by the nonlinear coupling between the concentration equations and Poisson’s equation, and closed-form solutions are available only in restricted geometries or in linearized regimes. A classical example is the Poisson–Boltzmann reduction near planar membranes, used to derive the Debye length; see (17)–(18). For more complex three-dimensional geometries, asymptotic methods have also been developed. In [2], asymptotic analysis of the PNP equations was used to derive reduced descriptions of synaptic potentials in the head and neck of dendritic spines, considering two ionic species. More recently, [23] derived asymptotic expressions for the voltage and ionic profiles in 3D nanodomains with currents entering and exiting through narrow window channels, allowing several ionic species of varying valence. The resulting current–voltage relations were validated against numerical simulations on spheroidal domains. In general, analytical approximations are available only in special cases, and the study of solutions of the PNP system therefore mostly relies on numerical approximations.

### 4.6 Concluding remark

In summary, the simplified analysis shows that the severe time-step restriction associated with operator splitting is a numerical artifact of the split treatment of the PNP equations. A coupled implicit formulation removes this restriction in the reduced model and performs accordingly in computations for the full system [7, 9]. This makes simulations of electrodiffusion on physiologically relevant time scales much more feasible, while still leaving open the separate question of how large the time step can be without compromising accuracy.

## Notes

### Competing Interest Statement

The authors have declared no competing interest.

### Funding

- The Research Council of Norway: 355113, 322312, 360005

## References

[1] Courtney L Lopreore, Thomas M Bartol, Jay S Coggan, Daniel X Keller, Gina E Sosinsky, Mark H Ellisman, and Terrence J Sejnowski. Computational modeling of three-dimensional electrodiffusion in biological systems: application to the node of Ranvier. Biophysical Journal, 95(6):2624–2635, 2008.

[2] Thibault Lagache, Krishna Jayant, and Rafael Yuste. Electrodiffusion models of synaptic potentials in dendritic spines. Journal of Computational Neuroscience, 47(1):77–89, 2019.

[3] Jerzy J Jasielec. Electrodiffusion phenomena in neuroscience and the Nernst–Planck–Poisson equations. Electrochem, 2(2):197–215, 2021.

[4] Jurgis Pods, Johannes Schönke, and Peter Bastian. Electrodiffusion models of neurons and extracellular space using the Poisson-Nernst-Planck equations – numerical simulation of the intra- and extracellular potential for an axon model. Biophysical Journal, 105(1):242–254, 2013.

[5] Jurgis Pods. A comparison of computational models for the extracellular potential of neurons. Journal of Integrative Neuroscience, 16(1):19–32, 2017.

[6] Karoline H Jæger, Ena Ivanovic, Jan P Kucera, and Aslak Tveito. Nano-scale solution of the Poisson-Nernst-Planck (PNP) equations in a fraction of two neighboring cells reveals the magnitude of intercellular electrochemical waves. PLoS Computational Biology, 19(2):e1010895, 2023.

[7] Karoline H Jæger and Aslak Tveito. Electrodiffusion dynamics in the cardiomyocyte dyad at nano-scale resolution using the Poisson-Nernst-Planck (PNP) equations. PLoS Computational Biology, 21(6):e1013149, 2025.

[8] Jinn-Liang Liu and Bob Eisenberg. Numerical methods for a Poisson-Nernst-Planck-Fermi model of biological ion channels. Physical Review E, 92(1):012711, 2015.

[9] Karoline H Jaeger and Aslak Tveito. Accurate computation of ionic concentrations in the synaptic cleft requires the full Poisson–Nernst–Planck (PNP) equations. bioRxiv, 2026.

[10] Maria G Kurnikova, Rob D Coalson, Peter Graf, and Abraham Nitzan. A lattice relaxation algorithm for three-dimensional Poisson-Nernst-Planck theory with application to ion transport through the gramicidin A channel. Biophysical Journal, 76(2):642–656, 1999.

[11] Benzhuo Lu, Michael J Holst, J Andrew McCammon, and YongCheng Zhou. Poisson–Nernst–Planck equations for simulating biomolecular diffusion–reaction processes I: Finite element solutions. Journal of Computational Physics, 229(19):6979–6994, 2010.

[12] Qiong Zheng, Duan Chen, and Guo-Wei Wei. Second-order Poisson–Nernst–Planck solver for ion transport. Journal of Computational Physics, 230(13):5239–5262, 2011.

[13] Jingwei Hu and Xiaodong Huang. A fully discrete positivity-preserving and energy-dissipative finite difference scheme for Poisson–Nernst–Planck equations. Numerische Mathematik, 145(1):77–115, 2020.

[14] Karoline H Jæger and Aslak Tveito. Differential Equations for Studies in Computational Electrophysiology, volume 14 of Simula SpringerBriefs on Computing. Springer Nature, 2023.

[15] Peter J. Mohr, David B. Newell, Barry N. Taylor, and E. Tiesinga. NIST reference on constants, units, and uncertainty. https://physics.nist.gov/cuu/Constants/index.html, Fundamental Constants Data Center of the NIST Physical Measurement Laboratory, 2018. Accessed: 2022-04-03.

[16] Andreas Solbrå, Aslak Wigdahl Bergersen, Jonas van den Brink, Anders Malthe-Sørenssen, Gaute T Einevoll, and Geir Halnes. A Kirchhoff-Nernst-Planck framework for modeling large scale extracellular electrodiffusion surrounding morphologically detailed neurons. PLoS Computational Biology, 14(10):e1006510, 2018.

[17] Dmitri A Rusakov, Leonid P Savtchenko, Kaiyu Zheng, and Jeremy M Henley. Shaping the synaptic signal: molecular mobility inside and outside the cleft. Trends in Neurosciences, 34(7):359–369, 2011.

[18] H Cohen and JW Cooley. The numerical solution of the time-dependent nernst-planck equations. Biophysical Journal, 5(2):145–162, 1965.

[19] Yoichiro Mori and Charles Peskin. A numerical method for cellular electrophysiology based on the electrodiffusion equations with internal boundary conditions at membranes. Communications in Applied Mathematics and Computational Science, 4(1):85–134, 2009.

[20] Edmund JF Dickinson, Juan G Limon-Petersen, and Richard G Compton. The electroneutrality approximation in electrochemistry. Journal of Solid State Electrochemistry, 15(7):1335–1345, 2011.

[21] Ning Qian and TJ Sejnowski. An electro-diffusion model for computing membrane potentials and ionic concentrations in branching dendrites, spines and axons. Biological Cybernetics, 62(1):1–15, 1989.

[22] A Peskoff and DM Bers. Electrodiffusion of ions approaching the mouth of a conducting membrane channel. Biophysical Journal, 53(6):863–875, 1988.

[23] F Paquin-Lefebvre, A Barea Moreno, and D Holcman. Voltage laws in nanodomains revealed by asymptotics and numerical simulations of electrodiffusion equations. Multiscale Modeling & Simulation, 24(1):365–397, 2026.

